# Efficacy of Biofilm Disrupters Against *Candida auris* and Other *Candida* species

**DOI:** 10.1101/2020.12.02.409250

**Authors:** Claudia A Cox, Jose A Vazquez, Sushama Wakade, Marek Bogacz, Matthew Myntti, Elias K Manavathu

## Abstract

**Background:** *C. auris* has become a globally emerging fungal pathogen, frequently reported to be multi-drug resistant, commonly found with *Staphylococcus aureus* in polymicrobial nosocomial infections. Although chlorhexidine (CHD) has been shown to be effective, it is associated with serious anaphylaxis reactions. Biofilm disrupters (BDs) are novel agents with a broad spectrum of antimicrobial activity. BDs have been used in the management of chronic wounds and to sterilize environmental surfaces. The goal of this study was to evaluate BDs against polymicrobial biofilms compared to CHD.

**Methodology:** We evaluated various BDs (BlastX, Torrent, NSSD) and CHD against *Candida spp* and *S. aureus* polymicrobial biofilms by zone of inhibition, biofilm, and time-kill assays. Effectiveness was based on the inhibition zone and the reduction of CFU, respectively, compared to the drug-free control.

**Results:** All BDs and CHD inhibited *C. auris* growth effectively in a concentration-dependent manner. Additionally, CHD and the BDs all showed excellent antimicrobial activity against polymicrobial biofilms. BDs were all highly effective against both *C. auris* isolates, whereas CHD was only moderately effective against *C. auris* 0386, suggesting resistance/tolerance. A comparative analysis of the BDs and CHD against *C. auris and C. albicans* by biofilm kill-curves showed at least 99.999% killing.

**Conclusions:** All three BDs and CHD have excellent activity against different *Candida* species, including *C. auris*. However, certain isolates of *C. auris* showed resistance/tolerance to CHD, but not to the BDs. The fungicidal activity of these novel agents will be valuable in eradicating surface colonization of *Candida spp*, including *C. auris*.

## INTRODUCTION

*Candida auris* is an emerging multidrug-resistant fungal pathogen that presents a serious global health problem (1–3). First isolated in Japan in 2009, geographically distinct clades of *C. auris* have been classified worldwide in Europe, Asia, Africa, North and South America (1–5). Since the rapid global emergence of this pathogen, a review of *Candida* strains deposited in the SENTRY Antifungal Surveillance Program dating back to 2004 revealed only four misidentified *C. auris* cases, indicating this species as a new pathogen (5). Interestingly, of the 16 first reported cases of *C. auris* infection, all specimens were isolated from the patients’ ears (6–9), though this fungal pathogen has now been isolated from several body sites ranging from asymptomatic cutaneous colonization to bloodstream infections (6). In fact, in the first *C. auris* outbreak which took place over the course of 16 months in a healthcare setting in Europe, of the 50 patients who became colonized, 44% required antifungal treatment and 18% developed bloodstream infections despite infection prevention measures implemented after the first few cases had been identified (10).

*C. auris* infections largely originate from healthcare facilities (2–4), but are known to be transmitted from person-to-person due to the ability of *C. auris* to colonize the skin (6, 11, 12). In this way, infection prevention and control must be administered much in the way of methicillin-resistant *Staphylococcus aureus* (MRSA) and carbapenem-resistant *Enterobacteriacea* (CRE) (6). This is in stark contrast to other *Candida* species, where cases are attributed to a patient’s own microbiome leading to autoinfection (6). This novel yeast poses a serious threat because of its level of multidrug resistance (MDR) and how readily this *Candida* spp. spreads in the hospital setting (1–4, 11). In fact, once it has established itself in the hospital environment, it has been difficult to eradicate *C. auris* from hospital surfaces (1–3). Elimination of this MDR pathogen from a London care center that involved 34 patients in 2016 brought with it a cost of over GB£1 million (approximately US$1.3 million) for initial control and GB£58,000 per month (approximately US$83,000) the following year (13).

In addition, it is now well described that *C. auris* generates a very significant biofilm (1–4) that increases resistance to antifungal agents and decreases the efficacy of surface agent (3). In fact, biofilms have been found in 90% of catheter-associated infections, which is indicative of the incredible impact *Candida* can have in a hospital setting where catheter infections are the leading cause of morbidity and mortality (14).

In clinical settings *C. auris* is not the only threat, as *Candida spp.* are the most common fungal pathogen identified in mycotic infections (15). Though *Candida spp.* are found commensally and normally do not cause disease, they are opportunistic and can cause infection (16, 17). While the causative agent behind most candidiasis cases has been *Candida albicans*, attributing to almost 50% of candidemia cases in the United States (18, 19) with a significant mortality rate of up to 47% in adults and 29% in children (20), other species have increasingly become clinically relevant (21). In fact, of at least 15 species able to infect humans, more than 90% of invasive cases are caused by only five: *C. albicans*, *C. glabrata*, *C. tropicalis*, *C. parapsilosis*, and *C. krusei* (21–23). Moreover, while most candidiasis are attributed to nosocomial acquisition, environmental acquisition is now described in the US and other developed countries (18).

It has been estimated that between 2002-2012, 22% of all invasive candidiasis cases requiring hospitalization in the US resulted in patient death with an average 21-day inpatient cost upwards of $45,000 (24). In a study to determine the annual national cost of fungal disease in the United States, the Centers for Disease Control and Prevention (CDC) found that hospitalizations resulting from *Candida* infections were estimated at $1.4 billion for 26,735 patients in 2017 alone (25).

Further, *Candida* colonizes several sites on the human body also inhabited by *Staphylococcus aureus*, a virulent, pathogenic, gram-positive bacteria (26). Co-infections with these microbes have been associated with higher mortality rates than those infections consisting of only one (26, 27). This is due, in part, to a synergistic relationship between the two microorganisms. This type of microbial relationship has created a significant need for broad-spectrum antimicrobial agents.

Chlorhexadine (CHD) has been used as a skin antiseptic for over half a century, and is effective against a wide range of microbes: gram-positive and gram-negative bacteria, yeast, and even enveloped viruses like HIV (28). However, CHD has also been linked to hypersensitivity reactions, up to and including anaphylaxis (29), and while anaphylactic reactions are rare there has been a rise in reported cases (30). Because of its wide spectrum of antimicrobial activity and low cost, it is unknowingly omnipresent in products extending beyond its disinfectant properties serving to possibly increase sensitization leading to a possible rise in allergic reactions to the product (29). To that end, it is necessary to find non-toxic alternative disinfectants.

The most commonly used antifungal agents are used systemically to treat patients with disseminated or invasive fungal infections. Unfortunately, *C. auris* is MDR, and thus is tolerant or resistant to many conventional antifungal agents. Moreover, environmental surfaces are generally cleaned with surface disinfectants such as CHD and other quaternary ammonium compounds. Recently, a non-toxic novel topical agent has been developed that has a unique mechanism of action. This new product is a high-osmolarity surfactant solution that has shown *in vitro* activity against gram-positive cocci, gram-negative bacilli, fungi and yeast (31, 32). Products that use this technology are also known as biofilm disrupting agents (BDs). These products are composed of a surfactant (benzalkonium chloride) and a high osmolarity buffer at a pH of 4.0 (33). In essence, the highly concentrated acid component destroys the extracellular polymeric substance (EPS) or biofilm by removing ionic metal bonds between EPS polymers and allowing for the penetration of the high-osmolarity solution which destroys bacterial and fungal cells. In addition, the pH is maintained for a prolonged period of time (~ 5 days) and also affects the persister cells so they can’t re-develop new biofilm.

The primary objective of this study is to evaluate the efficacy of three novel BDs formulated as a topical gel (BlastX), wound wash (Torrent), and a surface disinfectant (NSSD) and compare their activity to CHD on *C. auris, C. albicans, and C. glabrata in vitro* and in monomicrobial and polymicrobial biofilms composed of either *C. auris* or *C. albicans/S. aureus.*

## RESULTS

### All microbicidal agents tested effectively inhibit *Candida* growth

In order to determine the ability of each of the antimicrobial effects of BlastX, Torrent, and NSSD, in comparison to those of CHD and Puracyn, two of the leading commercial microbicides, inhibition-zone assays were conducted on *Candida albicans* 90028, *Candida auris* AR0381, and *Candida glabrata* 200918 biofilms using sterile saline as a negative control. This involved seeding Sabouraud’s Dextrose (SD) agar plates with *Candida* cells and then small volumes of each experimental compound to determine the effect on growth and biofilm formation and then measuring the zones of inhibition. All compounds were effective in inhibiting the growth of *Candida* cells, resulting in clear zones of inhibition (Figs. 1 and 2). Commercially available Puracyn, however, exhibited the smallest diameters of clearance against all strains and isolates when compared to all other tested compounds (Fig. 2).

**FIG 1.**
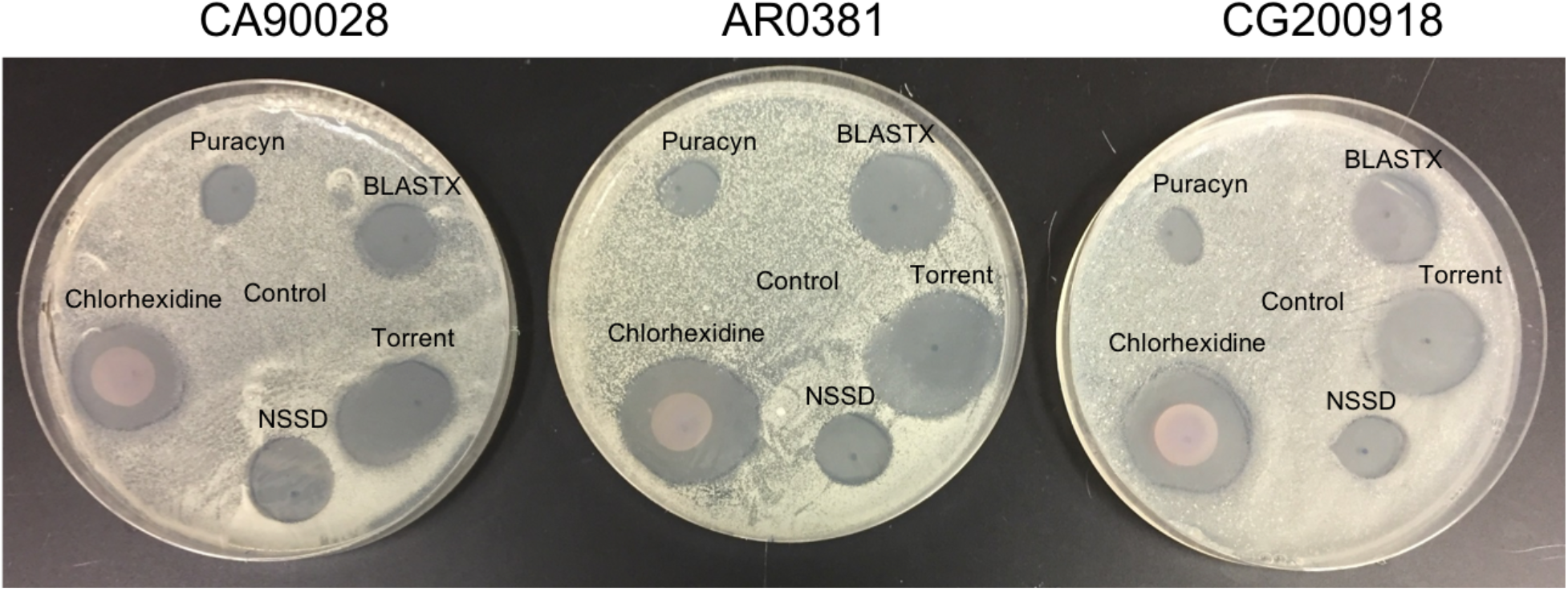
Photographic images showing the effects of various experimental and leading commercially available microbicidal agents on the growth and biofilm formation of *Candida albicans* 90028, *Candida auris* AR0381 and *Candida glabrata* 200918 using Sabouraud’s Dextrose (SD) agar plate inhibition assay. Briefly, SD agar plates were seeded with various *Candida* cells (1 × 10^5^ cells/plate) and the plates were air-dried under a biosafety cabinet to remove excess water. A uniform amount (20 μl) of the full-strength microbicidal agents (except BlastX , 50%) were applied strategically to the SD agar plates as shown. The plates were incubated at 37°C for 48 h in a plastic sleeve for microbial growth to obtain a clear zone of inhibition. Equal volume of sterile saline was used as control.

**FIG 2.**
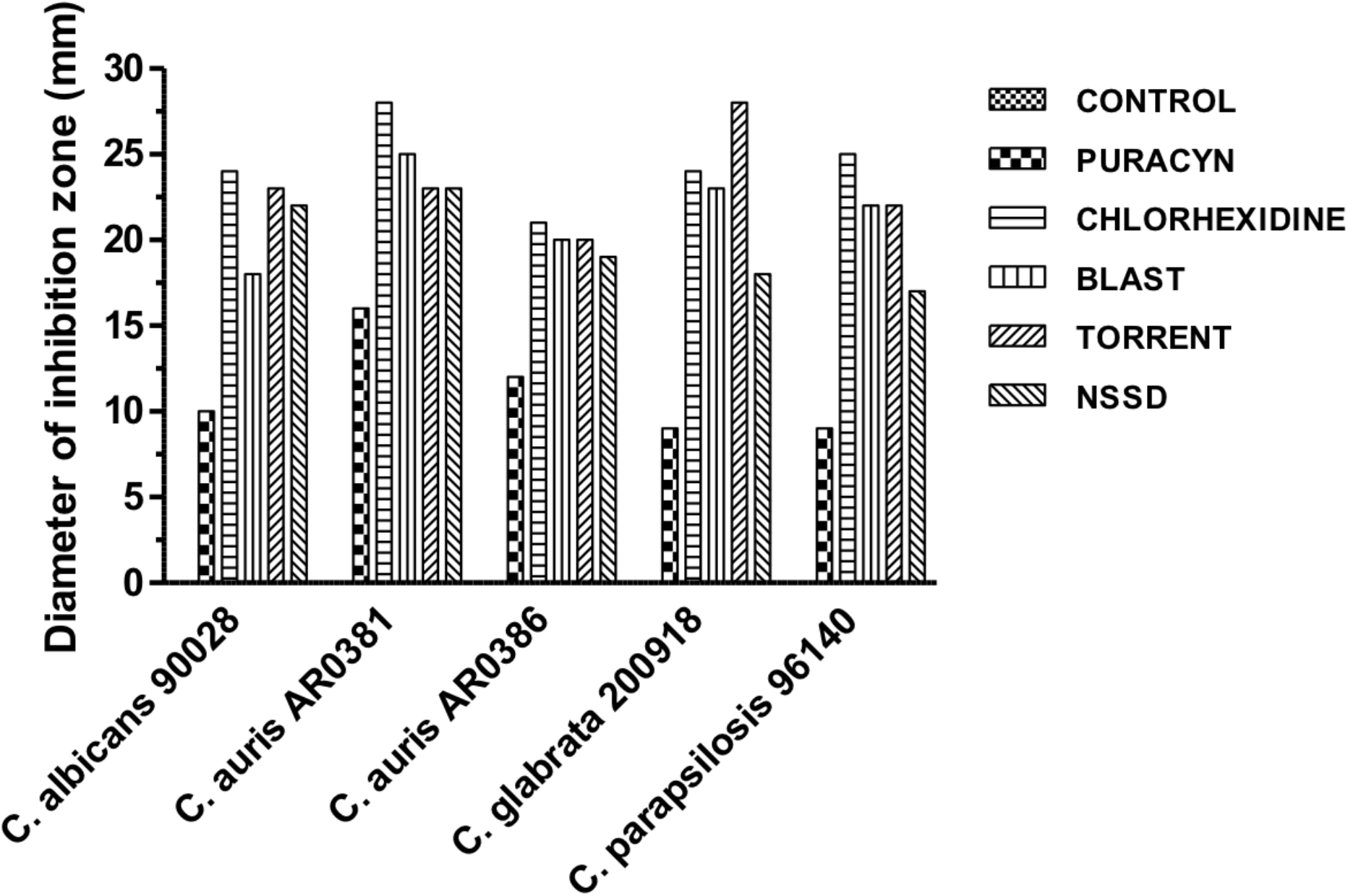
Effects of experimental and leading commercially available microbicidal agents on the growth and biofilm formation of various *Candida* species on Sabouraud’s Dextrose (SD) agar. SD agar plates were seeded with 1 × 10^5^ *Candida* cells/plate and a uniform amount (40 μl) of various microbicidal agents was applied to the SD agar and the plates were incubated at 37°C for 48 h for microbial growth and the development of clear zone of inhibition. The diameters of the inhibition zone obtained for various microbicidal agents were measured for each *Candida* species using a ruler and plotted. All microbicidal agents were used at full-strength except BlastX (50%).

### Monomicrobial fungal and bacterial biofilms are eliminated by all experimental microbicidal agents and CHD

To examine the effect of the BDs against monomicrobial biofilms in aqueous medium, *C. albicans*, *C. auris* 0381, *C. auris* 0386, and *S. aureus* biofilms were grown in 96-well plates. These biofilms were then treated with each agent for 24 hours and plated for CFU to determine the efficacy of each antiseptic. Puracyn was again seen to be the least effective among the tested agents, while all others were similarly effective against each of the organisms included (Fig. 3). As such, it was determined to cease testing of this compound in subsequent experiments.

**FIG 3.**
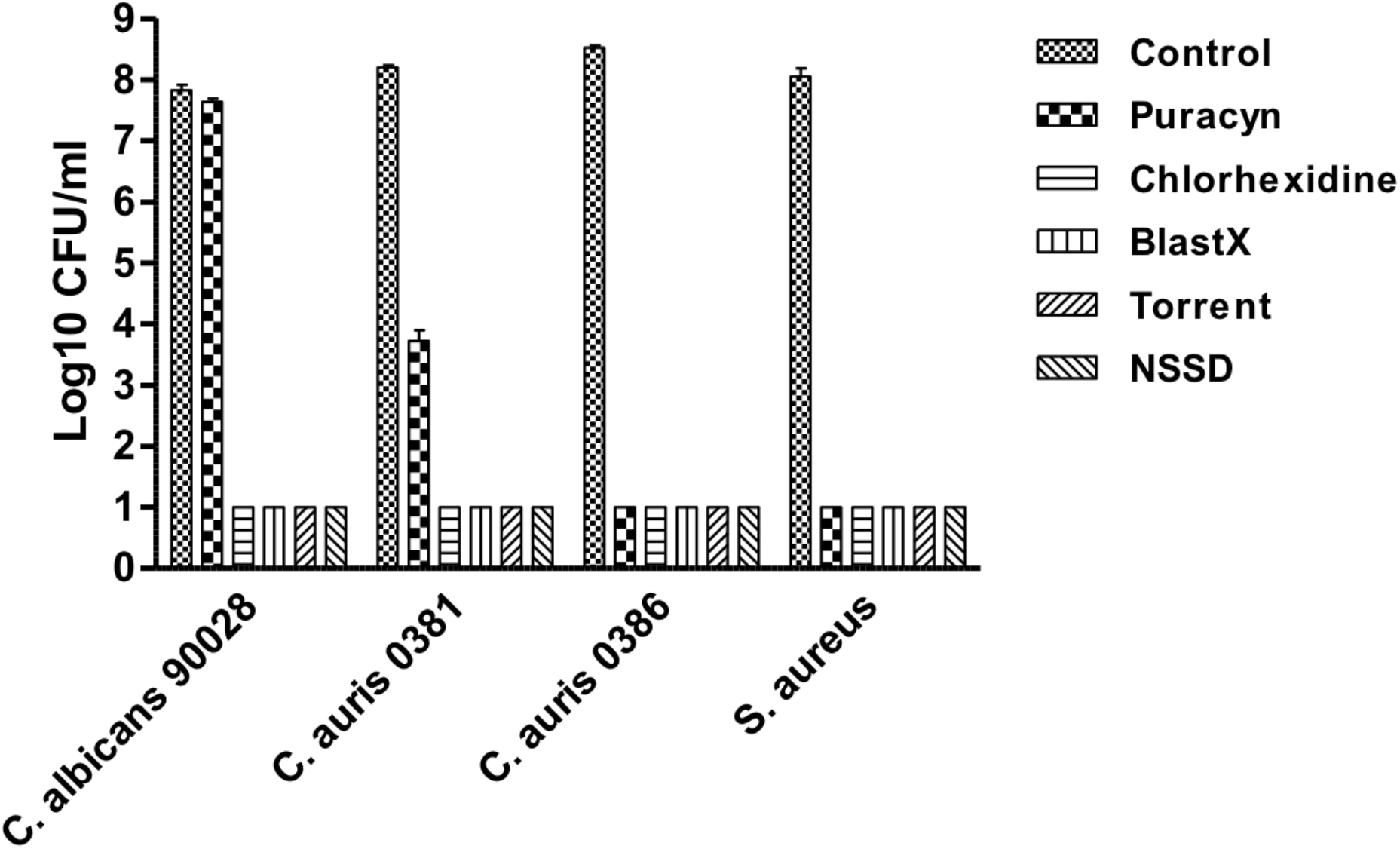
Effects of experimental and commercially available microbicides on candidal and bacterial monomicrobial biofilms. *C. albicans* 90028, *C. auris* 0381, *C. auris* 0386 and *S. aureus* 43300 were used in this study. Monomicrobial biofilms were developed in 96-well microtiter plates for 24 h at 37°C by incubating 0.1 ml cell suspension containing 1 × 10^7^ cells/ml in either Sabouraud’s dextrose broth (*Candida*) or Brain Heart Infusion broth (bacteria) as previously described. The biofilms were washed and exposed to full-strength experimental and commercial microbicides (except BlastX, 50%) for 24 h. The microbicide-treated biofilms were washed and their effectiveness was assessed by biofilm CFU assay. The control represents biofilm treated identically with PBS. The threshold of detection for the CFU assay was a minimum of 10. The experiment was repeated once and similar results were obtained.

### Time-kill assays indicate NSSD effectively eliminates *Candida auris*

As CHD has been associated with anaphylactic shock and toxicity, it was important to determine whether the microbicidal properties of the formulation of the experimental BDs was effective at eliminating *Candida* cells in a timely and dose-dependent manner. To ascertain this, time kill assays of suspensions of 10^7^/mL *C. albicans*, *C. glabrata*, and *C. auris* 0381 cells were treated with full-strength or 20% concentrations (in SD broth) of either NSSD or CHD for different time intervals, washed, and then plated to determine survival after treatment by counting CFU. At the 20% concentration, NSSD was slower to eradicate *Candida* cells than CHD (Fig. 4 A-C). However, at full-strength NSSD effectively killed all *Candida* species in under one minute, as did CHD (Fig. 4 D-F).

**FIG 4.**
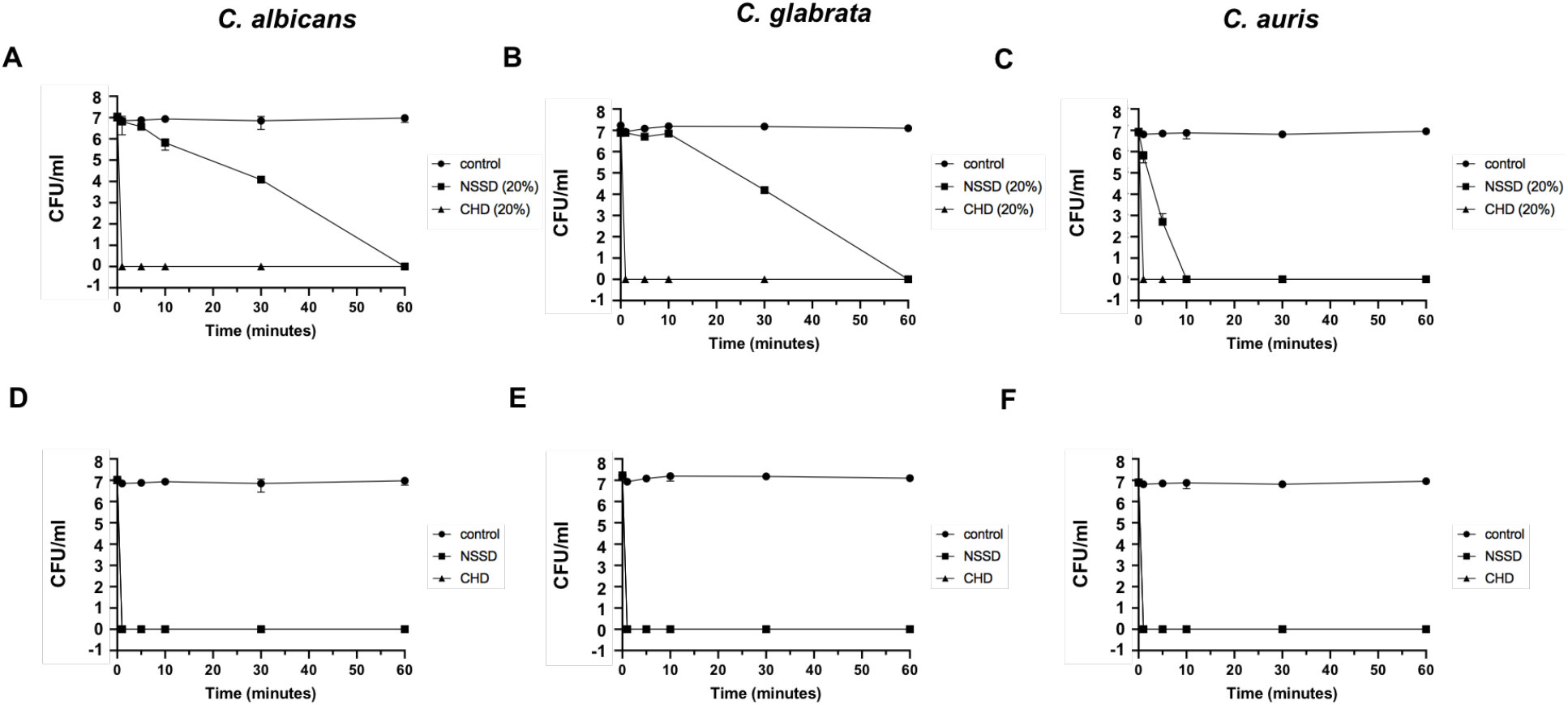
Time kill assays showing the effect of Next Science surface disinfectant and 4% chlorhexidine solution. *C. albicans* 90028 (A, D), *C. glabrata* 200918 (B, E), and *C. auris* 0381 (C, F) were used in this study. Cells were washed, pelleted, and resuspended in 20% solution (in SD broth) or full strength NSSD or CHD to a suspension of 1× 10^7^ cells/ml. At 1, 5, 10, 30, and 60 minute time intervals, cells were removed and washed, at which point serial dilutions were made and plated on SD agar plates. After 48 hours of incubation at 35°C, CFU were counted. This experiment was performed twice with similar results.

### Polymicrobial biofilms grown for 24 and 48 hours yield significant fungal and bacterial CFU

Most biofilms found environmentally are polymicrobial and as *C. albicans* is often co-cultured with *S. aureus* in bloodstream infections (17) and those infections related to biofilms (27), it was necessary to evaluate the experimental BDs in a similar model. In order to determine the optimal conditions in which to test polymicrobial biofilms consisting of different *Candida* species/isolates with *S. aureus*, biofilms were grown in 96-well, flat-bottomed plates seeded with 1 × 10^6^ *Candida* cells and 2 × 10^6^ *Staphylococcus* cells which were grown for 24 and 48 hours and counted for CFU. Both 24- (Fig. 5 A and B) and 48-hour (Fig. 5 C and D) time points produced robust biofilm, as evidenced by the high CFU yield, however the 48-hour samples generated a log_10_ higher CFU count. In an effort to produce more readily countable results, the 24-hour conditions were chosen to test the experimental compounds in subsequent experiments.

**FIG 5.**
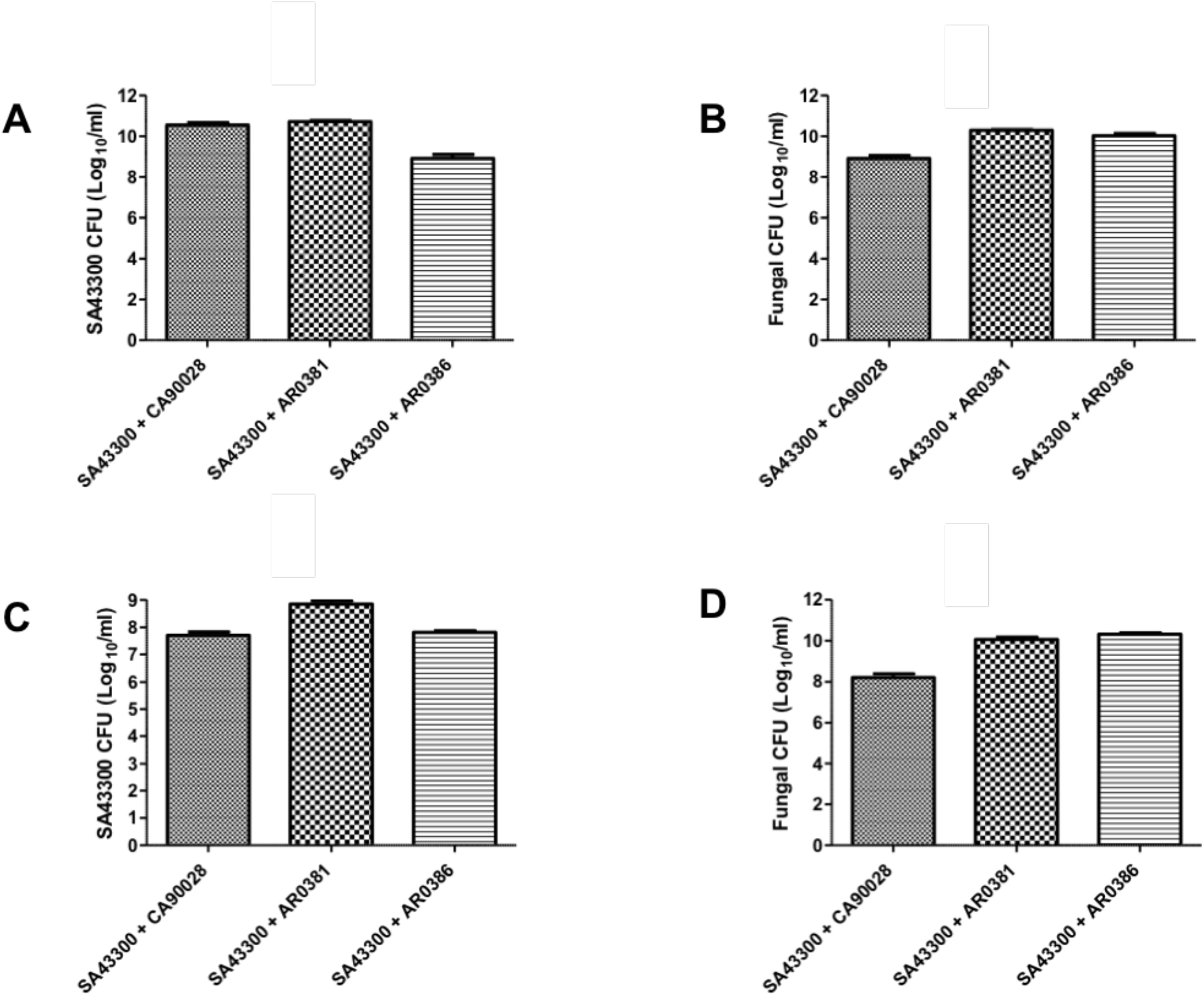
Twenty-four vs forty-eight hour *Candida* and *Staphylococcus aureus* polymicrobial biofilm. Dual species polymicrobial biofilms of *C. albicans* 90028, *C. auris* AR0381 and *C. auris* AR0386 with *S. aureus* 43300 were developed as previously described (Larkin *et al* AAC 61: e02396-16, 2017). Briefly, 0.1 ml suspensions of *Candida* (1 × 10^7^ cells/ml) were incubated in 50% fetal calf serum in 96-well flat bottom TC plates at 37°C for 90 min for cell adhesion. After cell adhesion, the plates were gently shaken on a gyratory shaker for 5 min and the unbound cells were removed with a pipette. The attached *Candida* cells were then incubated with 0.2 ml SA43300 cell suspension (1 × 10^7^ cells/ml) in SD broth for 24 or 48 h for biofilm development. The biofilms were washed with sterile distilled water two times and then the bacterial and the fungal CFUs were determined.

**FIG 6.**
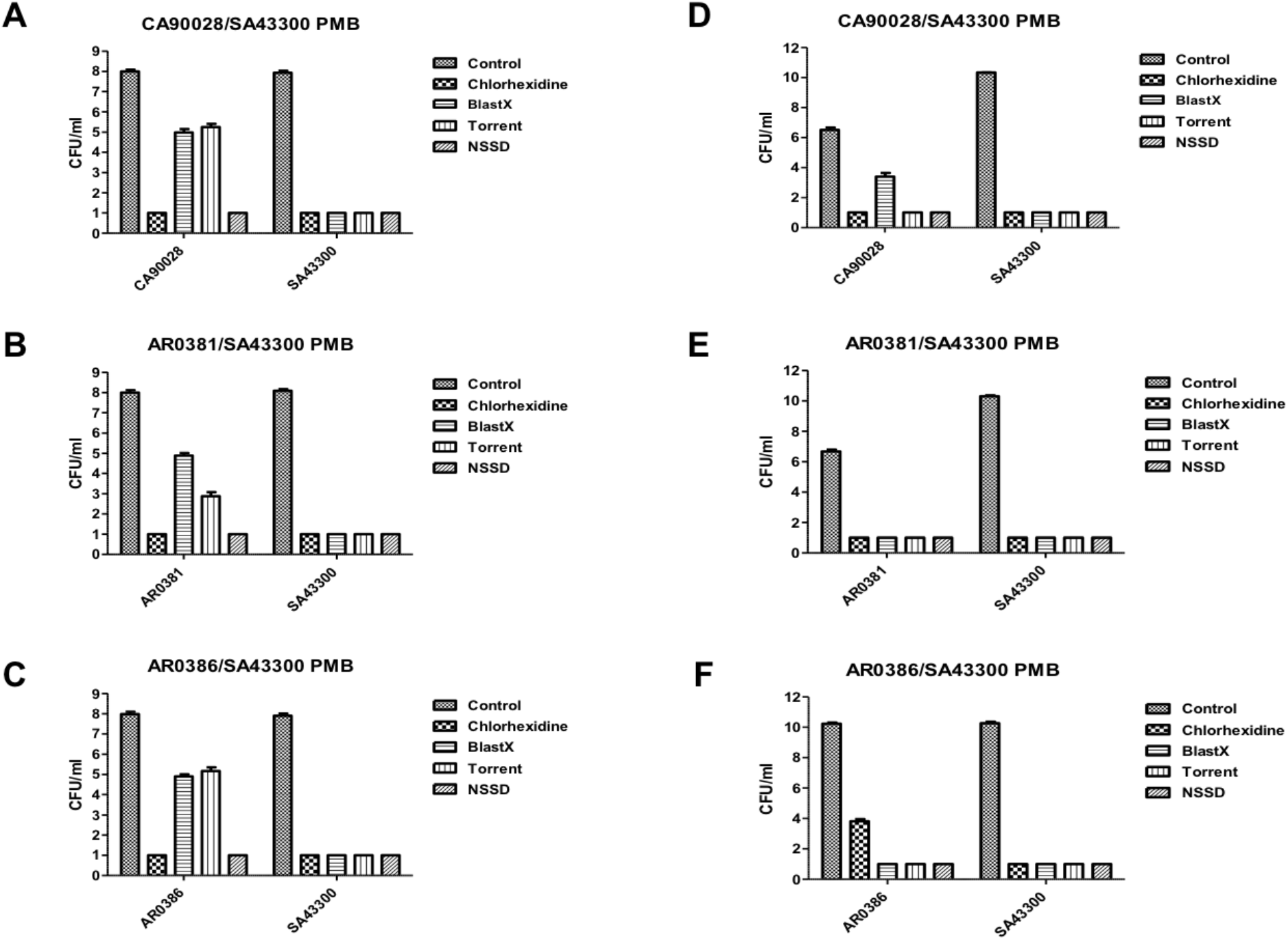
Antibiofilm activities of commercial and experimental microbicides. Dual species polymicrobial biofilms of *C. albicans* 90028, *C. auris* AR0381 and *C. auris* AR0386 with *S. aureus* 43300 were developed as previously described (Larkin et al AAC 61: e02396-16, 2017) in 96-well (A-C) or 24-well (D-F) TC plates for 24 h at 37°C. The biofilms were washed with sterile distilled water two times and then treated with various microbicidal agents prepared in SD broth at a concentration of 50% (A-C) or full strength (D-F), except BlastX which was 50%, for 24 h at 37°C. The microbicide-treated biofilms were washed with sterile distilled water three times, resuspended in sterile water and the fungal and bacterial CFUs were determined.

### Polymicrobial biofilms of *Candida* species and *S. aureus* are effectively killed by NSSD

This testing was done to determine the efficacy of each of the commercial and experimental antimicrobial agents in treating polymicrobial biofilms. To that effect, dual species biofilms of *C. albicans*, *C. auris* AR0381, and *C. auris* AR0386 with *S. aureus* were grown for 24 hours in 96-well plates. These biofilms were then treated for 24 hours with either 50% dilution or full-strength microbicidal agents (except Blast X, which was used consistently at 50% concentration) and plated for CFU. Though all BDs and CHD were effective against the biofilms, Torrent was unable to eliminate all fungal biofilm growth, yielding only ≥ 99.9% of fungal cells at 50% concentration, as well as BlastX, consistently tested at 50% concentration, failing to kill all fungal cells of all three tested *Candida* species on a similar scale to Torrent. Full-strength CHD did not fully kill all *C. auris* 0386 cells, yielding≥ 99.9% of fungal cell killing. The NSSD was able to fully kill all *C. auris* strains at both 50% and 100% concentration.

## DISCUSSION

*C. auris* is an emerging fungus which clearly has many aspects in which it is clinically relevant. Not only in finding ways in which to treat established infections, but also to curb the infectious aspect of this MDR pathogen. Most recently, efforts to extend the use of personal protective equipment (PPE) due to the COVID-19 pandemic has been acknowledged as a possible contributing factor to a surge in *C. auris* infections in southern California during the 2^nd^ and 3^rd^ quarters of 2020 (34). This provides additional support on the increasing importance of finding additional non-specific antiseptics to manage and inhibit outbreaks such as this.

In this study, we investigated the microbicidal activity of three different biofilm disrupters (BDs): a topical gel (BlastX), a wound wash solution (Torrent), and a topical disinfectant (NSSD); and compared them against commercially available 4% chlorhexidine (CHD) and Puracyn. These antimicrobial compounds were all tested against mono- and polymicrobial biofilms consisting of *C. auris*, *C. albicans*, *C. glabrata*, and *S. aureus* by zone of inhibition, time-kill, and biofilm assays.

All tested compounds were effective at inhibiting the growth of *Candida* species biofilms formed on SD agar plates, however Puracyn was least effective though this product has been shown to be bactericidal against MRSA (35). All three BDs and CHD inhibited the growth of polymicrobial biofilms with *C. auris* AR0381 and AR0386 in a concentration-dependent manner. However, full-strength CHD was less effective against *C. auris* AR0386 when compared to the full-strength experimental BDs. These biofilm experiments show that both the BDs and CHD are fungicidal against *C. auris* at a low concentrations. Furthermore, all three BDs also showed excellent activity against monomicrobial and polymicrobial biofilms consisting of *C. auris/S. aureus*. This is highly encouraging, as it is extremely important to identify antiseptic compounds with broad-spectrum activity, as polymicrobial biofilms are inherently difficult to eradicate from medical devices (26).

*S. aureus* polymicrobial biofilms have been shown to have increased resistance to several antibiotics (vancomycin, doxycycline, nafcillin, and oxacillin) (26). Add to that the fact that both *S. aureus* and *Candida* species contain several multidrug efflux pumps (1, 5, 6, 12, 14, 21, 26). This highlights the need for a non-specific antimicrobial disinfectant. Moreover, it has been suggested that repeated use of CHD at sub-inhibitory concentrations can lead to reduced activity against several oral pathogens. This may be due to the development of multidrug efflux pumps which can be expressed in both gram-positive and gram-negative bacteria (36).

Though highly effective against several species of *Candida* (37), the broad-spectrum antiseptic CHD has been shown to be less effective against *C. auris*, which can persist even after twice a day treatment with the antiseptic (38). One study showed that in clinically-relevant doses, mature biofilms were actually resistant to CHD treatment when compared to early biofilms and in a dose-dependent manner (39). To wit, there is a necessity for products designed to disrupt the extracellular polymeric substance of the biofilm, but is also effective against persister cells.

In conclusion, these novel BDs demonstrate excellent broad spectrum antimicrobial and antifungal activity which makes them very valuable in eradicating environmental surfaces and open wounds that may have been colonized with *Candida* species, including the MDR-*C. auris*, and thus possibly decrease the environmental spread of this deadly *Candida* species.

## MATERIALS AND METHODS

### Microorganisms and routine growth conditions

*Candida auris* AR0381, AR0386 (Mycology Section, Center for Disease Control), *Candida albicans* 90028 and *Candida glabrata 200918* (ATCC), and *S. aureus* 43300 (ATCC) were used in this study. Glycerol stocks of *Candida* species were stored at −80°C, cultured on SD agar plates, and incubated at 37°C. *S. aureus* was

### Agents Evaluated

Three biofilm disrupting agents (BDs) created by Next Science (Jacksonville, FL): a topical gel (BlastX), wound wash (Torrent), and a surface disinfectant (NSSD) against commercially available cleansers 4% chlorhexidine solution (CHD) and/or Puracyn Plus (Innovacyn Inc.).

### Sabouraud’s Dextrose (SD) agar zone inhibition assay

SD agar plates were seeded with 1 × 10^5^ *Candida* cells and air-dried in a laminar flow hood until excess liquid was absorbed. To these, 20 μl full-strength (Puracyn, CHD, Torrent, NSSD) or 50% (BlastX) commercial and experimental compounds or sterile saline solution (control) were carefully pipetted onto the surface of the agar. Plates were incubated at 37°C for 48 hours in a plastic sleeve. Diameters of the zones of inhibition were measured in mm by ruler.

### Time kill assays

*Candida spp*. cultures were grown overnight in SD broth at 35°C. Cells were washed by centrifugation at 2,000 x g for 10 minutes and were subsequently resuspended in 100% or 20% solution (in SD broth) of NSSD and CHD or SD broth at final density of 1 × 10^7^ cells/ml. At 1, 5, 10, 30, and 60 min time points, 100 μl of the cell suspension was removed and washed with 1 ml of brain heart infusion (BHI) broth. Serial dilutions in BHI broth were made and cells were seeded onto SD agar plates. CFU was assessed after 48 hours of incubation at 35°C. All experiments were performed twice.

### Monomicrobial biofilm assays

Biofilms were grown by seeding 96-well microtiter plates with 100 μl of a 1 × 10^7^ cells/ml suspension in either SD broth (*Candida*) or BHI broth (*Staphylococcus*) for 24 hours at 37°C. Biofilms were washed and treated with full-strength microbicidal compounds (except Blast X, 50%) or PBS (control) for 24 hours. Treated biofilms were then washed and viable cells counted by biofilm CFU assay, where the threshold of detection was a minimum of 10 colonies. Experiments were performed twice with similar results.

### Polymicrobial biofilm assays

Biofilms of *C. albicans* 90028, *C. auris* 0381, and *C. auris* 0386 with *S. aureus* 43300 were grown as previously described (40). Briefly, 500 or 100 μl suspensions of *Candida* (1 × 10^7^/ml cells) were seeded in 24- or 96-well plates in 50% fetal calf serum and incubated at 37°C for 90 minutes for cell adhesion. Plates were then gently shaken on a gyratory shaker for 5 minutes, after which planktonic cells were removed via pipette. The attached *Candida* cells were then incubated with 500 or 200 μl suspensions *S. aureus* suspension (1 × 10^7^/ml) in SD broth for 24 hours for biofilm formation. Biofilms were then washed with sterile water twice and treated with antimicrobial agents at full-strength or 50% concentration (except BlastX, always 50%) for 24 hours at 37°C. Biofilms were then washed with sterile water three times, resuspended in either 1 ml or 100 μl sterile water, at which point fungal and bacterial CFUs were determined.

## ACKNOWLEDGMENTS

Funding and biofilm disrupting solutions were provided by Next Science LLC, Jacksonville, FL.

## References

1. Bidaud AL, Chowdhary A, Dannaoui E. 2018. Candida auris: An emerging drug resistant yeast - A mini-review. J Mycol Med 28:568–573.

2. Sears D, Schwartz BS. 2017. Candida auris: An emerging multidrug-resistant pathogen. Int J Infect Dis 63:95–98.

3. Kean R, Ramage G. 2019. Combined Antifungal Resistance and Biofilm Tolerance: the Global Threat of Candida auris. mSphere 4.

4. Semreen MH, Soliman SSM, Saeed BQ, Alqarihi A, Uppuluri P, Ibrahim AS. 2019. Metabolic Profiling of Candida auris, a Newly-Emerging Multi-Drug Resistant Candida Species, by GC-MS. Molecules 24.

5. Spivak ES, Hanson KE. 2018. Candida auris: an Emerging Fungal Pathogen. J Clin Microbiol 56.

6. Forsberg K, Woodworth K, Walters M, Berkow EL, Jackson B, Chiller T, Vallabhaneni S. 2019. Candida auris: The recent emergence of a multidrug-resistant fungal pathogen. Med Mycol 57:1–12.

7. Satoh K, Makimura K, Hasumi Y, Nishiyama Y, Uchida K, Yamaguchi H. 2009. Candida auris sp. nov., a novel ascomycetous yeast isolated from the external ear canal of an inpatient in a Japanese hospital. Microbiol Immunol 53:41–4.

8. Oh BJ, Shin JH, Kim MN, Sung H, Lee K, Joo MY, Shin MG, Suh SP, Ryang DW. 2011. Biofilm formation and genotyping of Candida haemulonii, Candida pseudohaemulonii, and a proposed new species (Candida auris) isolates from Korea. Med Mycol 49:98–102.

9. Kim MN, Shin JH, Sung H, Lee K, Kim EC, Ryoo N, Lee JS, Jung SI, Park KH, Kee SJ, Kim SH, Shin MG, Suh SP, Ryang DW. 2009. Candida haemulonii and closely related species at 5 university hospitals in Korea: identification, antifungal susceptibility, and clinical features. Clin Infect Dis 48:e57–61.

10. Schelenz S, Hagen F, Rhodes JL, Abdolrasouli A, Chowdhary A, Hall A, Ryan L, Shackleton J, Trimlett R, Meis JF, Armstrong-James D, Fisher MC. 2016. First hospital outbreak of the globally emerging Candida auris in a European hospital. Antimicrob Resist Infect Control 5:35.

11. Armstrong PA, Rivera SM, Escandon P, Caceres DH, Chow N, Stuckey MJ, Diaz J, Gomez A, Velez N, Espinosa-Bode A, Salcedo S, Marin A, Berrio I, Varon C, Guzman A, Perez-Franco JE, Escobar JD, Villalobos N, Correa JM, Litvintseva AP, Lockhart SR, Fagan R, Chiller TM, Jackson B, Pacheco O. 2019. Hospital-Associated Multicenter Outbreak of Emerging Fungus Candida auris, Colombia, 2016. Emerg Infect Dis 25.

12. de Jong AW, Hagen F. 2019. Attack, Defend and Persist: How the Fungal Pathogen Candida auris was Able to Emerge Globally in Healthcare Environments. Mycopathologia 184:353–365.

13. Taori SK, Khonyongwa K, Hayden I, Athukorala GDA, Letters A, Fife A, Desai N, Borman AM. 2019. Candida auris outbreak: Mortality, interventions and cost of sustaining control. J Infect 79:601–611.

14. Sardi JCO, Scorzoni L, Bernardi T, Fusco-Almeida AM, Mendes Giannini MJS. 2013. Candida species: current epidemiology, pathogenicity, biofilm formation, natural antifungal products and new therapeutic options. J Med Microbiol 62:10–24.

15. Deorukhkar SC, Saini S, Mathew S. 2014. Non-albicans Candida Infection: An Emerging Threat. Interdiscip Perspect Infect Dis 2014:615958.

16. Vazquez-Gonzalez D, Perusquia-Ortiz AM, Hundeiker M, Bonifaz A. 2013. Opportunistic yeast infections: candidiasis, cryptococcosis, trichosporonosis and geotrichosis. J Dtsch Dermatol Ges 11:381–93; quiz 394.

17. Carolus H, Van Dyck K, Van Dijck P. 2019. Candida albicans and Staphylococcus Species: A Threatening Twosome. Front Microbiol 10:2162.

18. Pappas PG, Lionakis MS, Arendrup MC, Ostrosky-Zeichner L, Kullberg BJ. 2018. Invasive candidiasis. Nat Rev Dis Primers 4:18026.

19. Kullberg BJ, Arendrup MC. 2015. Invasive Candidiasis. N Engl J Med 373:1445–56.

20. Pappas PG, Rex JH, Lee J, Hamill RJ, Larsen RA, Powderly W, Kauffman CA, Hyslop N, Mangino JE, Chapman S, Horowitz HW, Edwards JE, Dismukes WE, Group NMS. 2003. A prospective observational study of candidemia: epidemiology, therapy, and influences on mortality in hospitalized adult and pediatric patients. Clin Infect Dis 37:634–43.

21. Sanguinetti M, Posteraro B, Lass-Florl C. 2015. Antifungal drug resistance among Candida species: mechanisms and clinical impact. Mycoses 58 Suppl 2:2–13.

22. Pfaller MA, Messer SA, Moet GJ, Jones RN, Castanheira M. 2011. Candida bloodstream infections: comparison of species distribution and resistance to echinocandin and azole antifungal agents in Intensive Care Unit (ICU) and non-ICU settings in the SENTRY Antimicrobial Surveillance Program (2008-2009). Int J Antimicrob Agents 38:65–9.

23. Pappas PG, Kauffman CA, Andes DR, Clancy CJ, Marr KA, Ostrosky-Zeichner L, Reboli AC, Schuster MG, Vazquez JA, Walsh TJ, Zaoutis TE, Sobel JD. 2016. Clinical Practice Guideline for the Management of Candidiasis: 2016 Update by the Infectious Diseases Society of America. Clin Infect Dis 62:e1–50.

24. Strollo S, Lionakis MS, Adjemian J, Steiner CA, Prevots DR. 2016. Epidemiology of Hospitalizations Associated with Invasive Candidiasis, United States, 2002-2012(1). Emerg Infect Dis 23:7–13.

25. Benedict K, Jackson BR, Chiller T, Beer KD. 2019. Estimation of Direct Healthcare Costs of Fungal Diseases in the United States. Clin Infect Dis 68:1791–1797.

26. Todd OA, Peters BM. 2019. Candida albicans and Staphylococcus aureus Pathogenicity and Polymicrobial Interactions: Lessons beyond Koch’s Postulates. J Fungi (Basel) 5.

27. Schlecht LM, Peters BM, Krom BP, Freiberg JA, Hansch GM, Filler SG, Jabra-Rizk MA, Shirtliff ME. 2015. Systemic Staphylococcus aureus infection mediated by Candida albicans hyphal invasion of mucosal tissue. Microbiology (Reading) 161:168–181.

28. Edmiston CE, Jr., Bruden B, Rucinski MC, Henen C, Graham MB, Lewis BL. 2013. Reducing the risk of surgical site infections: does chlorhexidine gluconate provide a risk reduction benefit? Am J Infect Control 41:S49–55.

29. Moka E, Argyra E, Siafaka I, Vadalouca A. 2015. Chlorhexidine: Hypersensitivity and anaphylactic reactions in the perioperative setting. J Anaesthesiol Clin Pharmacol 31:145–8.

30. Fernandes M, Lourenco T, Lopes A, Spinola Santos A, Pereira Santos MC, Pereira Barbosa M. 2019. Chlorhexidine: a hidden life-threatening allergen. Asia Pac Allergy 9:e29.

31. Miller KG, Tran PL, Haley CL, Kruzek C, Colmer-Hamood JA, Myntti M, Hamood AN. 2014. Next science wound gel technology, a novel agent that inhibits biofilm development by gram-positive and gram-negative wound pathogens. Antimicrob Agents Chemother 58:3060–72.

32. Banerjee D, Tran PL, Colmer-Hamood JA, Wang JC, Myntti M, Cordero J, Hamood AN. 2015. The antimicrobial agent, Next-Science, inhibits the development of Staphylococcus aureus and Pseudomonas aeruginosa biofilms on tympanostomy tubes. Int J Pediatr Otorhinolaryngol 79:1909–14.

33. Wolcott R. 2015. Disrupting the biofilm matrix improves wound healing outcomes. J Wound Care 24:366–71.

34. Health CDoP. 2020. Health Advisory: Resurgence of Candida auris in Healthcare Facilities in the Setting of COVID-19. State of California-Health and Human Services Agency, http://publichealth.lacounty.gov/eprp/lahan/alerts/CAHANCauris082020.pdf.

35. Rani SA, Hoon R, Najafi RR, Khosrovi B, Wang L, Debabov D. 2014. The in vitro antimicrobial activity of wound and skin cleansers at nontoxic concentrations. Adv Skin Wound Care 27:65–9.

36. Cieplik F, Jakubovics NS, Buchalla W, Maisch T, Hellwig E, Al-Ahmad A. 2019. Resistance Toward Chlorhexidine in Oral Bacteria - Is There Cause for Concern? Front Microbiol 10:587.

37. Salim N, Moore C, Silikas N, Satterthwaite J, Rautemaa R. 2013. Chlorhexidine is a highly effective topical broad-spectrum agent against Candida spp. Int J Antimicrob Agents 41:65–9.

38. Ku TSN, Walraven CJ, Lee SA. 2018. Candida auris: Disinfectants and Implications for Infection Control. Front Microbiol 9:726.

39. Kean R, McKloud E, Townsend EM, Sherry L, Delaney C, Jones BL, Williams C, Ramage G. 2018. The comparative efficacy of antiseptics against Candida auris biofilms. Int J Antimicrob Agents 52:673–677.

40. Larkin E, Hager C, Chandra J, Mukherjee PK, Retuerto M, Salem I, Long L, Isham N, Kovanda L, Borroto-Esoda K, Wring S, Angulo D, Ghannoum M. 2017. The Emerging Pathogen Candida auris: Growth Phenotype, Virulence Factors, Activity of Antifungals, and Effect of SCY-078, a Novel Glucan Synthesis Inhibitor, on Growth Morphology and Biofilm Formation. Antimicrob Agents Chemother 61.

